# Sex Lability and Dimorphism in Dioecious Palmer Amaranth (*Amaranthus palmeri*)

**DOI:** 10.1101/769935

**Authors:** Mohsen B. Mesgaran, Maor Matzrafi, Sara Ohadi

## Abstract

Dioecious weeds (separate sexes) may benefit from a maximized outcrossing and optimal sex-specific resource allocation but there are costs associated with the evolution of this breeding system which can be exploited for long-term management of dioecious weeds. That is, seed production in dioecious species is contingent upon the co-occurrence and co-flowering of the two genders and can be further complicated by biases in sex ratio. We therefore explored the sex ratio and dimorphism in secondary sex characters in three populations of Palmer amaranth (*Amaranthus palmeri*) from California, Kansas and Texas and tested if water stress can change the sex expression and/or the synchrony of flowering (anthesis) between male and female plants. Sex ratio (proportion of males) was balanced and did not deviate from 1:1 in all experiments and populations (California, Kansas, Texas) when plants received normal watering. Male and female plants of *A. palmeri* did not differ in timing of emergence, plant height and relative growth rate. While the initiation of flowering (emergence of inflorescence) occurred earlier in males than females, females preceded males in timing of anthesis. Water stress delayed anthesis in males to a larger extent than females giving rise to an anthesis mismatch as large as seven days between the two sexes. Water stress induced a female sex expression in Kansas population giving rise to a female to male ratio of 1.78 which significantly differed from the equal 1:1 sex ratio. Our data provide the first evidence of sex lability and environment sex determination (ESD) in *A. palmeri*, suggesting manipulation of sex expression and phenological asynchrony as novel tools for ecological management of dioecious weeds.

## Introduction

Hermaphroditism represents the most common mode of sexual reproduction in weeds whilst dioecy (separate male and female individuals) is rare and have been observed in about 5-7% of weed species (Sutherland 2004). Whereas a selfing breeding system is often associated with a weedy character (Baker 1955; Razanajatovo et al. 2016), it might come as a surprise that some dioecious weeds such as palmer amaranth (*Amaranthus palmeri* S. Watson) and common waterhemp (*A. tuberculatus* (Moq.) J. D. Sauer) have been reported amongst the most the troublesome weeds in US (Van Wychen 2017). Contrary to hermaphroditism (particularly selfing), dioecy maximizes outcrossing and thereby reduces the likelihood of inbreeding depression (Thomson and Barrett 1981,1990; Charlesworth 1999). Dioecy also has the benefit of optimizing the allocation of resources between male and female functions by minimizing the competition between these two functions through sexual specialization, i.e. the so-call “division of labor” theory (Maynard Smith 1978; Charnov 1982).

Despite these advantages, a dioecious breeding system is subject to several demographic handicaps. First, the two sexes must co-occur in the same habitat and, unlike hermaphrodite weeds, dioecious plants cannot propagate from a single individual. Second, the two sexes need to flower at the same time. The smaller the degree of temporal overlapping between male and female flowering, the lower the chance of successful fertilization and hence seed production. Third, the fecundity of a dioecious population can be affected by the sex ratio (the proportion of males) and is likely to decline if the population becomes skewed towards one sex: a male-biased population may not achieve its maximum seed production because of the ovule limitation, whilst in a female-biased population, limited pollen availability may reduce the seed output. Therefore, it seems that there exists an optimal sex ratio at which the population-level seed production is maximized, but beyond that optimal point, the seed output is likely to decline due to pollen or ovule limitations (Mulcahy 1967).

We believe that novel weed management strategies can be developed by better understanding of the above “cracks” in the evolution of a dioicous mating system. Two potential strategies envisioned involve shifting the sex ratio towards one sex (sub-optimal sex ratio strategy) and reducing the flowering overlap between the two sexes (phenological isolation strategy). The proposed strategies rely on the assumptions that the sex is labile–hence sex ratio can be changed–and that we can manipulate the flowering phenology of male and female individuals differentially. For long sex was posited to be a fixed trait of the plant throughout its life cycle and genetic sex determination (GSD), where genes/chromosomes specify the gender of an individual, was considered as the sole mechanism by which the sex of an individual (dioecious) plant is determined (Fuller and Ritchie 1967). Evidence, however, has mounted over the past half century showing that sex can be labile and the genetic control of sex can be overridden by the environment (reviewed in Freeman et al. 1980; Korpelainen 1998; Vega-Frutis et al. 2014): Environmental sex determination (ESD) has been found in many plants where the expression of sex has changed in response to the environmental factors such as soil moisture (Freeman and Vitale 1985), nutrients (Dorken and Barrett 2004), light intensity (Zimmerman 1991), photoperiod (Thompson 1956), plant density (Onyekwelu and Harper 1979), and exogenous application of plant hormones (Chailakhyan 1979). Environmental stresses such as water stress (drought) are more likely to induce maleness, while conditions conducive to growth e.g. high soil nutrition favor a female sexuality (Freeman et al., 1980), perhaps because reproduction is more costly to females than males (Delph, 1999; Meagher, 1988).

In many dioicous species, male and female individuals have shown to differ in secondary sex characters (i.e. traits other than the sex itself) related to morphology, physiology, life history and phenology (reviewed in Geber et al. 1999; Juvany and Munné-Bosc 2015; Lloyd and Webb 1977). Differences in progression of phenological events (e.g. timing of germination) can result in temporal segregation of males and females, ultimately reducing the chance of successful cross-pollination in a dioecious population. In fact, most dioecious species seems to be protandrous i.e. males flower earlier than females (Forero-Montaña and Zimmerman 2010; Forest 2014). As female’s investment in reproduction is greater than males, it has been hypothesized that females postpone flowering (reproduction) for longer period of resource accumulation (Lloyd and Webb 1977; Purrington and Schmitt 1998). This hypothesis has direct implications for manipulative separation of flowering timing between the two sexes. That is, if requirement for resource accumulation delays flowering in females, then environmental stresses can further isolate the females, exacerbating the flowering mismatch between the two sexes.

In this study, we attempted to test the viability of exploiting the demographic handicap associated with a unisexual breeding system (i.e. shift in sex ratio and phenological isolation) for novel management of dioecious weeds. We used palmer amaranth (*Amaranthus palmeri* S. Watson) as our dioecious model system. The species has been ranked as the worst weed in US corn fields in a recent survey (Van Wychen 2017) and evolved resistance to six classes of herbicides with 56 herbicide-resistant biotypes found in the US (Heap 2018). Most of these biotypes were found to be resistant to the widely used herbicide, glyphosate (Roundup®), requiring the development of non-chemical control methods. The objectives of this study were threefold: 1) determine if the male and female individuals of *A. palmeri* differ in secondary sex characters related to growth and development, 2) examine sex lability (shift in sex ratio) in *A. palmeri* in response to environmental stress (drought), and 3) test if the degree of flowering synchrony between *A. palmeri* male and female plants can be affected by the environmental stress (drought). As a proof of concept study, we used water stress for the sex change experiment as this environmental factor has been found to affect sex expression in several plant species (Freeman et al., 1980; Korpelainen, 1998).

## Materials and Methods

Seeds of *A. palmeri* from California (CA) and Kansas (KS) were cordially provided by Dr. Anil Shrestha (California State University, Fresno, California) and Dr. Dallas E. Peterson (Kansas State University, Manhattan, Kansas), respectively. Seeds of a third population were collected from mature plants (about 30 plants) from a set-aside field in College Station, Texas (TX), in the summer of 2017. Seeds were stored at 4°C before use. Two sets of experiments were conducted in the greenhouse facility (Orchard Parks) of University of California, Davis, during the spring and summer of 2018.

### Experiment 1. Natural sex ratio and dimorphism in secondary sex characters

The aim of this experiment was to examine whether *A. palmeri* has undergone divergent evolution in secondary sex traits related to growth and development. We therefore compared male and female plants of *A. palmeri* for potential differences in emergence pattern, timing of flowering and anthesis, plant height, and relative growth rate (RGR). By using three populations (CA, KS, and TX) and large number of plants (∼250), *Experiment 1* also served as a platform for a robust estimate of baseline sex ratio in *A. palmeri*, which was the major trait to be examined in the second experiment (*Experiment 2*, see below). In July 2018, we sow 200 seeds from each population, into 5 by 5 by 5 cm plastic pots (one seed per pot) filled with a soil mix (1:1:1:3 sand:compost:peat:dolomite). We recorded the emergence time for each individual seed and then transplanted the young plants (two-four leaf stage) into larger plastic pots (2.37 L) filled with a soil mix (1:1 sand/peat) plus a controlled-release fertilizer (15–9–12, 150 g 75 L^−1^; Scotts Osmocote PLUS, Mississauga, ON). Pots were randomly placed in a net house under natural California-summer conditions and watered twice a day through an automated drip irrigation. Plant height was measured three times (11 July, 23 July, and 24 September) during the season. We inspected plants daily and an individual was recorded to be at the flowering stage when the tip of its inflorescence was visible either on the terminal or lateral buds. The onset of anthesis for a flowering plant was then determined when the stigma (in pistillate flowers) or dehisced anther (in staminate flower) become visible. At this stage, i.e. anthesis, we also recorded the sex of plants.

### Experiment 2. Sex expression and flowering synchrony in response to water stress

This experiment aimed at evaluating the effect of water stress on the sex ratio and flowering synchrony between the male and female plants of *A. palmeri*. Because of space limitation we only used two populations (CA and KS) in this experiment. The experiment was conducted twice (April-July and July-September in 2018) in a completely randomized design with 50 replicates (plants) per treatment (water stress *vs.* well-watered control). About 10 seeds were sown in each round plastic pot (2.37 L), filled with a soil mix (1:1 sand/peat) plus a controlled-release fertilizer (15 – 9 – 12, 150 g 75 L^−1^; Scotts Osmocote PLUS, Mississauga, ON). Pots were placed in a greenhouse set at a temperature of 32/22°C (day/night) with a day length of 16 hours provided through supplementary lighting between 5:00 to 9:00 am and 5:00 to 9:00 pm. Seedling were thinned haphazardly several times to achieve one plant per pot by their four-leaf stage.

When plants reached six to seven true-leaf stage, 50 random plants from each population were randomly assigned to well-watered or water deficit treatments. Well-watered plants received water from four emitters inserted into the potting medium to deliver 65 mL of water min^−1^ for two mins and twice per day (7 am and 2:00 pm). Water deficit treatment was achieved by removing three out of four drip emitters from designated water stress treatment pots, reducing the amount of water by approximately 75%. Additional irrigation (∼100 mL twice per week) was added to water deficit plants when severe visual stress symptoms were observed. Plants were sexed by examining flowers for the presence of anther (in males) or stigma (in females). For each plant we also measured time to flowering (inflorescences) and time to anthesis as described in *Experiment 1*.

### Statistical analysis

*In Experiment 1*, a binomial test (two-tailed) was used to test if the observed sex ratio in each population statistically deviates from the expected 1:1 sex ratio (i.e. 50% male). Binomial test was conducted by ‘binom.test()’ function in ‘stats’ package of R (R Core Team, 2018). Using a t-test, male and female plants were compared for the secondary sex characters including time to flowering (inflorescence), time to anthesis, plant height, and early stage RGR. Function ‘t.test()’ of ‘stats’ package (R Core Team, 2018) was used for the t-test analysis. The early stage RGR was calculated based on the height gain between the first and second height measurements using the following formula (Radosevich et al. 2007):

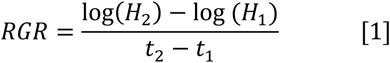

where *H_1_* and *H_2_* are plant heights measured at time *t_1_* (11 July) and *t_2_* (23 July), respectively (log stands for natural logarithm). To explore difference in seedling emergence pattern between the sexes, we fitted the following two-parameter logistic equation to cumulative emergence data of *A. palmeri* using the ‘drm()’ function of ‘drc’ package (Ritz et al. 2016):

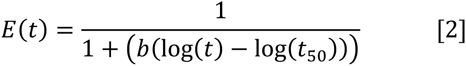

where *E(t)* is the cumulative emergence over time, *t*, *t_50_* is time to 50% emergence while *b* denotes the steepness of emergence curve around *t_50_*.

In *Experiment 2*, for each water treatment level we tested the deviation of sex ratio from 1:1 by using a binomial test (two-tailed) as described above. Phenology data (time to inflorescence and time to anthesis) were subject to analysis of variance, ANOVA, using the ‘lm()’ and ‘Anova()’ functions of ‘stats’ (R Core Team, 2018) and ‘car’ (Fox et al. 2019) packages in R, respectively. Inspection of residuals showed no predictable pattern when plotted against fitted values suggesting that the ANOVA’s assumption of homogeneity of variance has been met. The degree of flowering/anthesis overlap between the two sexes was visually inspected by fitting a kernel density smoother to time data (i.e. time to flowering and time to anthesis) using the ‘geom_density_ridges()’ function of ‘ggplot2’ R package (Wickham et al. 2019).

## Results and Discussion

### Experiment 1. Natural sex ratio and dimorphism in secondary sex characters

#### Sex Ratio

The recorded sex ratio was balanced in all three tested populations and did not deviate from the expected 1:1 male/female ratio as indicated by a binomial test (Figure 1). Although California population had slightly higher proportion of males (57%) than females, the departure from 50% sex ratio was not significant (P = 0.306) (Figure 1).

**Figure 1.**
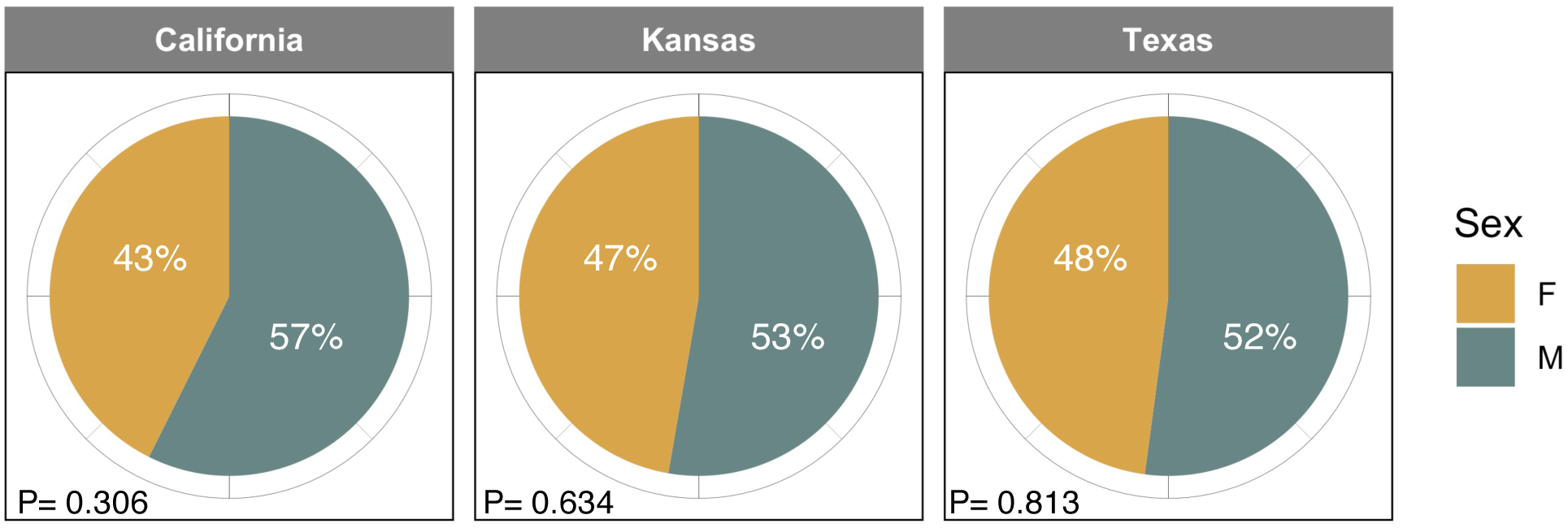
Observed sex ratios in three *A. palmeri* populations grown in pots during the summer of 2018 in California, Davis. Also reported are the P-values (shown as P) for testing the deviation of sex ratio (% male) from 50% based on a binomial test. Total number of plants examined for sex ratio estimation was 61, 110, and 76 for California, Kansas, and Texas. Abbreviations: F: Female and M: Male.

An equal sex ratio was found in *A. palmeri* plants grown under greenhouse conditions with no water or nutrient limitations (Giacomini et al., 2014). A balanced sex ratio of 55% (22 males vs. 18 females) also was reported by Lemen (1980); however, he does not provide any information about method of measuring this parameter e.g. whether sex has been determined in plants grown from the seed or sex ratio has been measured under field conditions. Keeley et al. (1987) also reported a male:female ratio of 47%:53%—which shows a balanced sex ratio—for a field grown *A. palmeri*. However, the source of seed for this test population was collected from one single plant suggesting that their reported sex ratio could best represent the sex ratio of that individual and not the population. Of 300 pot-grown *A. palmeri* plants—for a pollen mediate gene flow study—172 (∼60%) and 162 (%54) plants became male in the first (2005) and second year (2006) of experiment (Sosnoskie et al. 2012). The sex ratio has clearly (and statistically) been male-based in the first year according to a binomial test (*P-vale* = 0.0009), though the sex ratio data were not analyzed in the original article.

#### Seedling Emergence

Total emergence for California, Kansas, and Texas populations were 31%, 55%, and 35% (n=200 for each individual population), respectively. Logistic model (Eq.2) fitted the emergence data adequately as evidenced in Figure 2 and small root mean square errors (RMSE ≤ 4%) in all populations. Female and male plants of *A. palmeri* exhibited almost identical emergence pattern and in none of the tested populations the emergence curves differed between the sexes. Populations, however, differed in emergence rate (rapidity) with Kansas being the fastest emerging population (*t_50_* = 2.5±0.10 days) while California was the slowest (*t_50_* = 4.3±0.12 days) irrespective of the gender (see inset in Figure 2).

**Figure 2.**
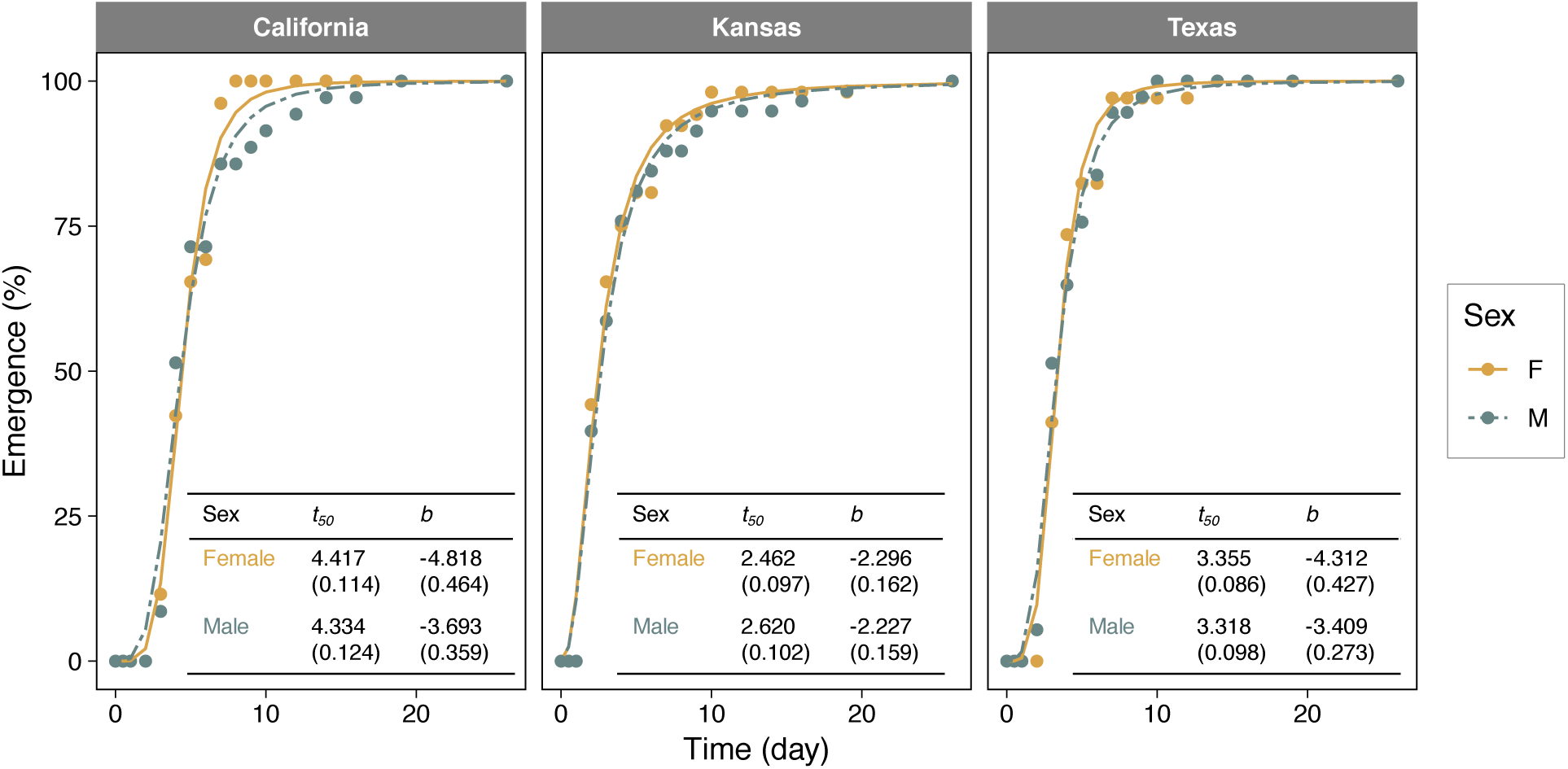
Comparison of cumulative seedling emergence between female (F) and male (M) plants in three populations *A. palmeri* originating. Inset tables show parameter estimates for the logistic model (equation 2). Values in the parentheses indicate stand errors. Note that emergence data were normalized relative to the final number of seedlings counted for each sex within the population (see text for definition of model’s parameters).

#### Time to Flowering and Anthesis

In all three populations, the initiation of flowering (i.e. tip of inflorescence becoming visible) occurred earlier in male than female plants of *A. palmeri* (Figure 3). The difference in timing of flowering was particularly profound in Texas population where males, on average, flowered ∼9 days before females. Variability in flowering time seemed to be similar between males and females as illustrated by the violin plots. The only exception was observed with California population where the flowering bulge occurred close to the earliest flowering times (i.e. at the bottom of the violin) as opposed to females that had their bulge above the median time of flowering. Note that a bulge on a violin plot represents the value (here flowering time) with the highest probability of occurrence. The male’s earlier flowering may suggest a protandrous flowering habit in *A. plameri*; however, when we investigated the timing of anthesis, which might be regard as the effective flowering event, we arrived at an opposite conclusion.

**Figure 3.**
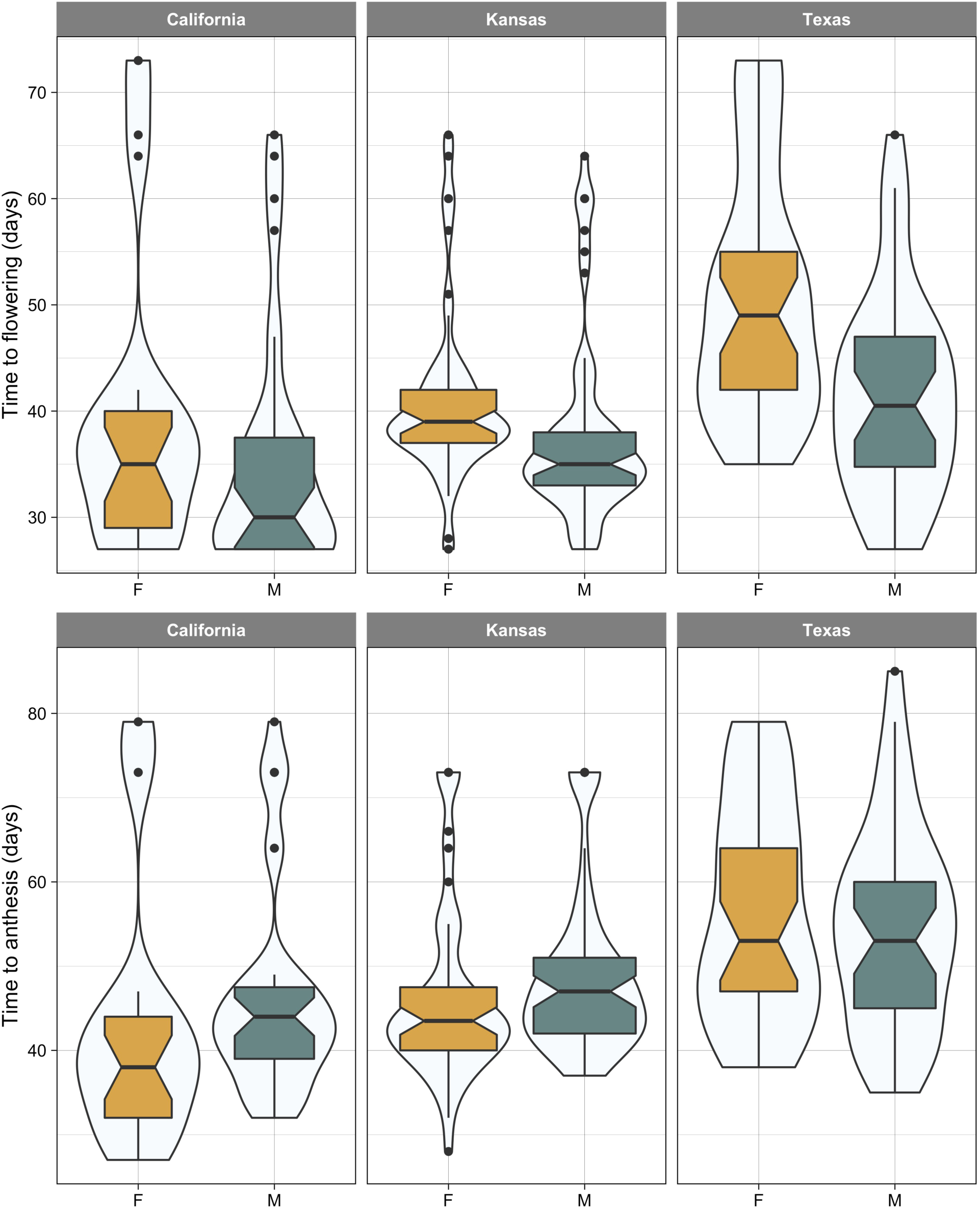
Days to first flowering (inflorescence visible; top row) and anthesis (bottom row) in female (F) and male (M) plants of *A. palmeri* from three populations (California, Kansas, and Texas). The width of violin plot represents the relative frequency of the event of interest (i.e. flowering or anthesis time). Horizontal line within the boxplot shows the median, the box includes the interquartile range, IQR, (i.e. 25^th^ to 75^th^ percentile range), whiskers extend 1.5 times the IQR, and filled circles represent extreme values which fall within > 1.5×IQR and < 3×IQR.

In both, California and Texas populations, anthesis happened four days earlier in females than males (Figure 3). For Texas population, although the median time to anthesis was the same for male and females, the most probable anthesis time (i.e. the bulge) was at an earlier time in females compared to males. While male plants of *A. palmeri* began to flower earlier than female plants, it took them a much longer time to mature as shown in Figure 4. For females, regardless of the population, the length of time from the appearance of inflorescence to anthesis (visible stigma) was no more than 4-5 days compared with a 11-day time period needed for males to develop mature anthers following the initiation of flowering. This observation suggests that *A. palmeri* should be classified as a protogynous species as its pistillate flowers mature earlier than the staminate ones (Figure 4). Korres et al. (2017) also showed that female plants of *A. palmeri* flowered 6 to 8 d earlier than male plants, a flowering pattern that is very similar to our result from Kansan and California population. However, Korres et al. (2017) do not explicitly define “flowering” stage and we are not sure if our “anthesis” represents the same reproductive stage as with the “flowering” stage in Korres et al. (2017). A synchronous anthesis between male and female plants observed with Texas population resembles the results of Giacomini et al. (2014) who found no differences in days to first flowering (anthesis) between male and female plants of *A. palmeri*.

**Figure 4.**
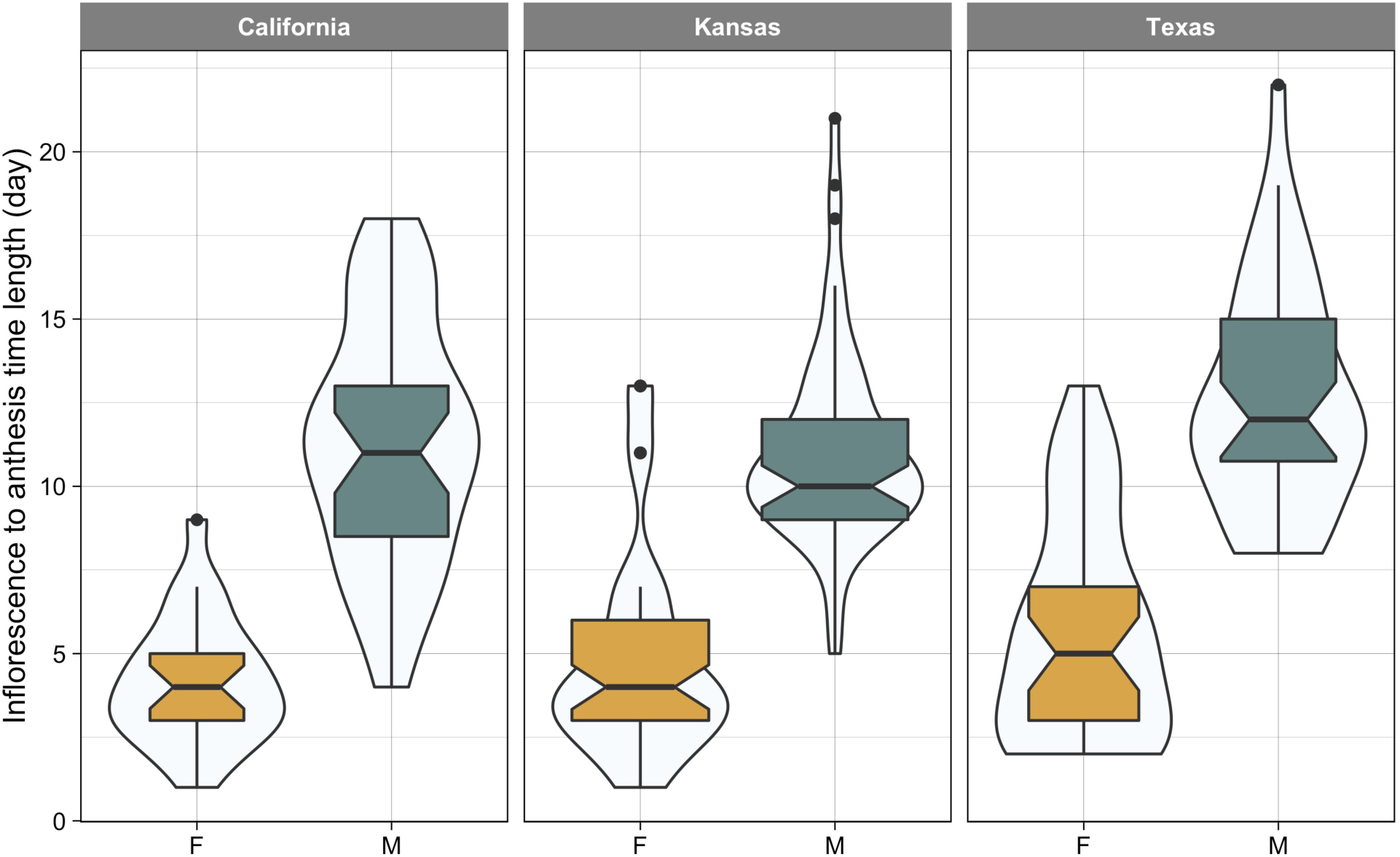
Length of time (day) from flowering (inflorescence visible) to anthesis in female (F) and male (M) plants of *A. palmeri* from three populations (California, Kansas, and Texas). The width of violin plot shows the relative frequency of event (inflorescence to anthesis time). The horizontal line within the boxplot shows the median, the box includes the interquartile range, IQR, (i.e. 25^th^ to 75^th^ percentile range), whiskers extend 1.5 times the IQR, and filled circles represent extreme values which fall within > 1.5×IQR and < 3×IQR.

#### Plant Height and Relative Growth Rate (RGR)

We observed no sex-specific differences in final plant height for the three tested populations (Figure 5). The only significant sexual dimorphism in growth was related to the height-based RGR in Texas population. At the early stage of growth, males of this population grew tall at a rate that was ∼13% faster than females. However, the male’s faster initial growth did not translate into significant larger plants as males attained a shorter stature (125.3 ± 3.78 cm) than females (133.3 ± 4.69 cm) at the end of growing season. In the study of Webster and Grey (2015) plant height did not differ between *A. palmeri* sexes either grown alone or in competition with cotton, but sex differential growth was detected in other traits e.g. females had wider canopy and produced more than double the biomass of males. Several other studies have shown that female *A. palmeri* plants tend to be taller and grow faster than males (e.g. Keeley et al., 1987; Korres et al., 2017). Whereas *A. palmeri* populations originating from ten contrasting cropping systems differed in many morphological and growth traits, no sexual dimorphism was reported in these populations (Bravo et al. 2017).

**Figure 5.**
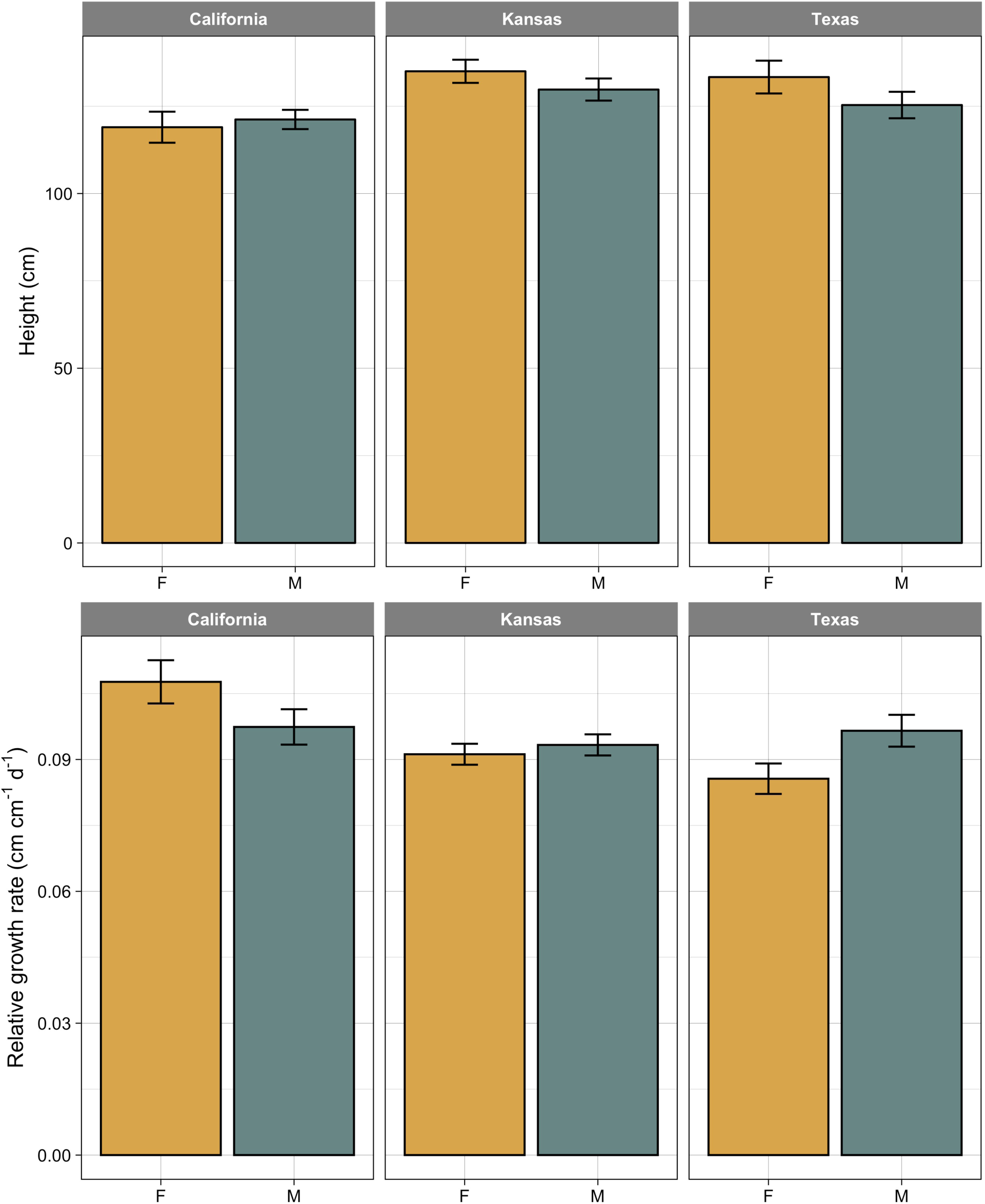
Compression of final plant height (top row) and relative growth rate (bottom row) between female (F) and male (M) plants in three populations of *A. palmeri*. Vertical lines on bars indicate 95% confidence interval.

### Experiment 2. Sex expression and flowering synchrony in response to water stress

#### Sex Ratio

While sex ratio remained balanced across water treatments in California population, exposure to drought produced a heavily female-biased sex ratio (∼1.78 times more female than males) in Kansas population. Sex ratio significantly deviated from 50% (P-value = 0.009, n = 99) in water stress treatment whilst it was balanced (P-value = 0.547, n = 99) when *A. palmeri* plants received normal irrigation (Figure 6). Our data provide the first evidence of sex lability and environment sex determination (ESD) in *A. palmeri:* water stress induced a female sex expression in Kansas population.

**Figure 6.**
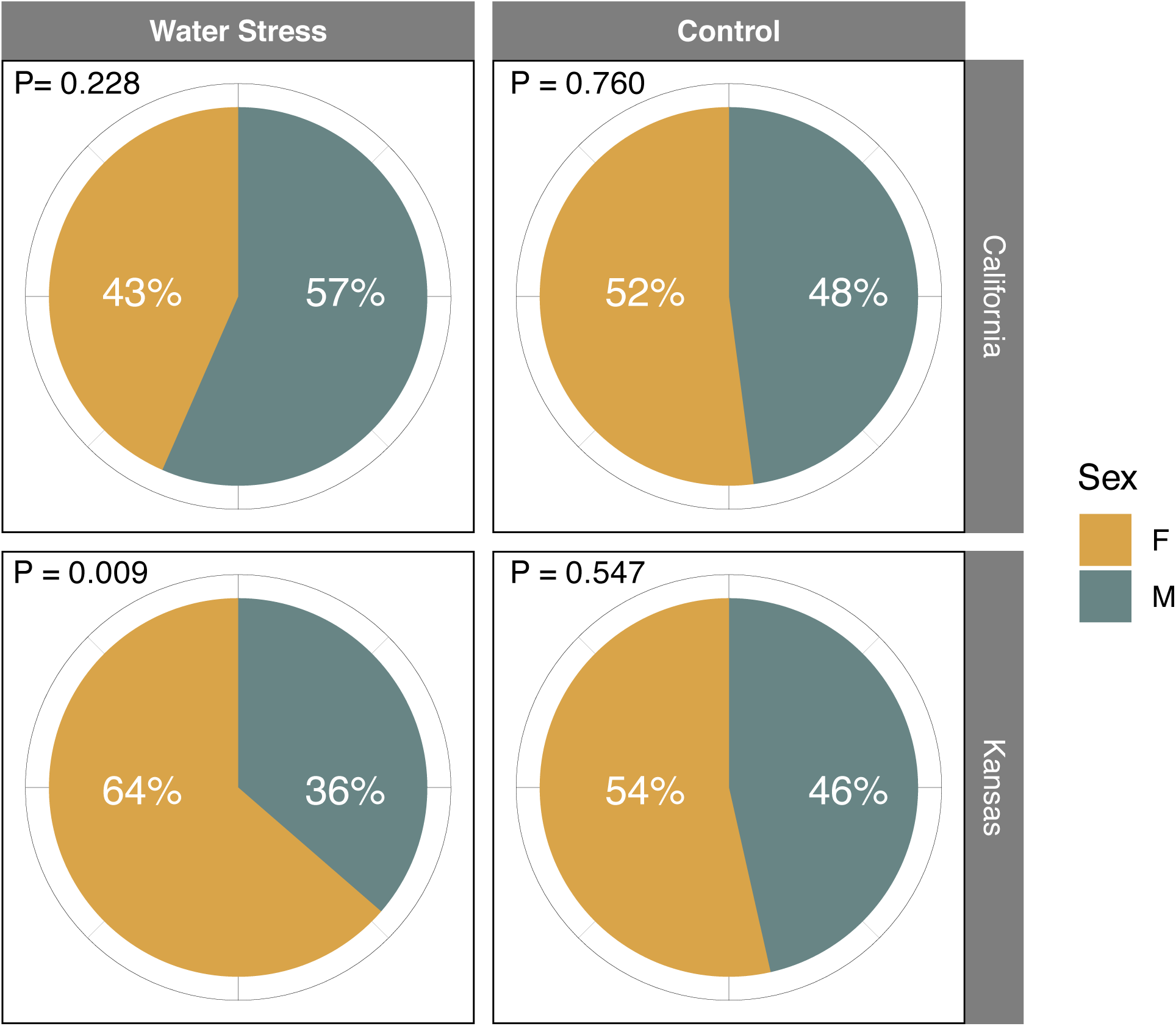
Observed sex ratios in two *A. palmeri* populations grown under well-watered (control) and water stress conditions. Also reported are the P-values (shown as P) for testing the deviation of sex ratio (% male) from 50% based on a binomial test. Total number of plants (sum of two experiments) examined for sex ratio estimation was ∼100 plants per water treatment. Abbreviations: F: Female and M: Male.

To the best of our knowledge, our study provides the first case of a feminizing effect for water stress: almost all previous work has reported a masculinizing effect for this stressor. Since reproduction is more costly to females than males (Delph 1999; Meagher 1988), sex allocation theory predicts that the environmental conditions conducive to plant growth should favor a female sex expression while resource-poor environments should induce maleness (Charnov 1982). Empirical data support the prediction of sex allocation theory. Reviews by Freeman et al. (1980) and Korpelainen (1998) show that water stress has induced maleness in all 11 species (five dioecious and six monoecious) that have been tested for potential sex change in response to draught. Limited field surveys have also recorded overrepresentation of males in resource-poor habitats whereas females were more abundant in benign habitats (Policansky, 1981; Lovett Doust and Cavers 1982). The observed overrepresentation of males under stressful environments may be due to a sex differential mortality rather than being the result of an actual sex change. Most studies on sex ratio, however, lack the capacity to disentangle these two processes i.e. sex lability *vs.* sex-differential survival. By randomly thinning seedlings (with no bias towards selection of vigorous seedlings), our study was able to rule out differential mortality as the potential driver of shift in sex ratio. In our experiments, only two plants (one per water treatment in Kansas population) died over the course of experiment.

#### Time to Flowering and Anthesis

Flowering phenology was recorded for plants from California and Kansas populations grown under different irrigation regimes. Similar to *Experiment 1*, the initiation of flowering happened earlier in males than females with water stress having no effects on the flowering sequence between the two sexes (Figure 7). However, water stress slightly delayed flowering, almost to the same extent, in both male and female plants giving rise to a more temporally staggered flowering pattern as compared with the control. Consistent with *Experiment 1*, *A. palmeri* female plants preceded males in timing of anthesis in the water stress experiment (*Experiment 2*; Figure 8). Sex-differential responses to water stress were observed with the timing of anthesis: water stress distorted the frequency distribution of anthesis timing to a larger extent in males than females (Figure 8). Time to anthesis nearly followed a unimodal distribution in both sexes when plants received normal irrigation but became multimodal and “wavy” when plants were exposed to water stress. Anthesis was much synchronized between male and female plants of *A. palmeri* under normal watering but the two sexes became more separate (i.e. less overlapping in anthesis) with water stress. In both populations, the difference between males and females in median time to anthesis (arrows in Figure 8) was only two days in control but extended to seven days in water-stressed plants. The increased asynchrony between the two sexes was mainly driven by a differential response of the sexes to water stress. While water stress delayed anthesis in both sexes, the delay in males was much longer than females (Figure 8). In California population, for example, water stress postponed anthesis (median time) by seven days in male plants compared with a 2-day delay in females.

**Figure 7.**
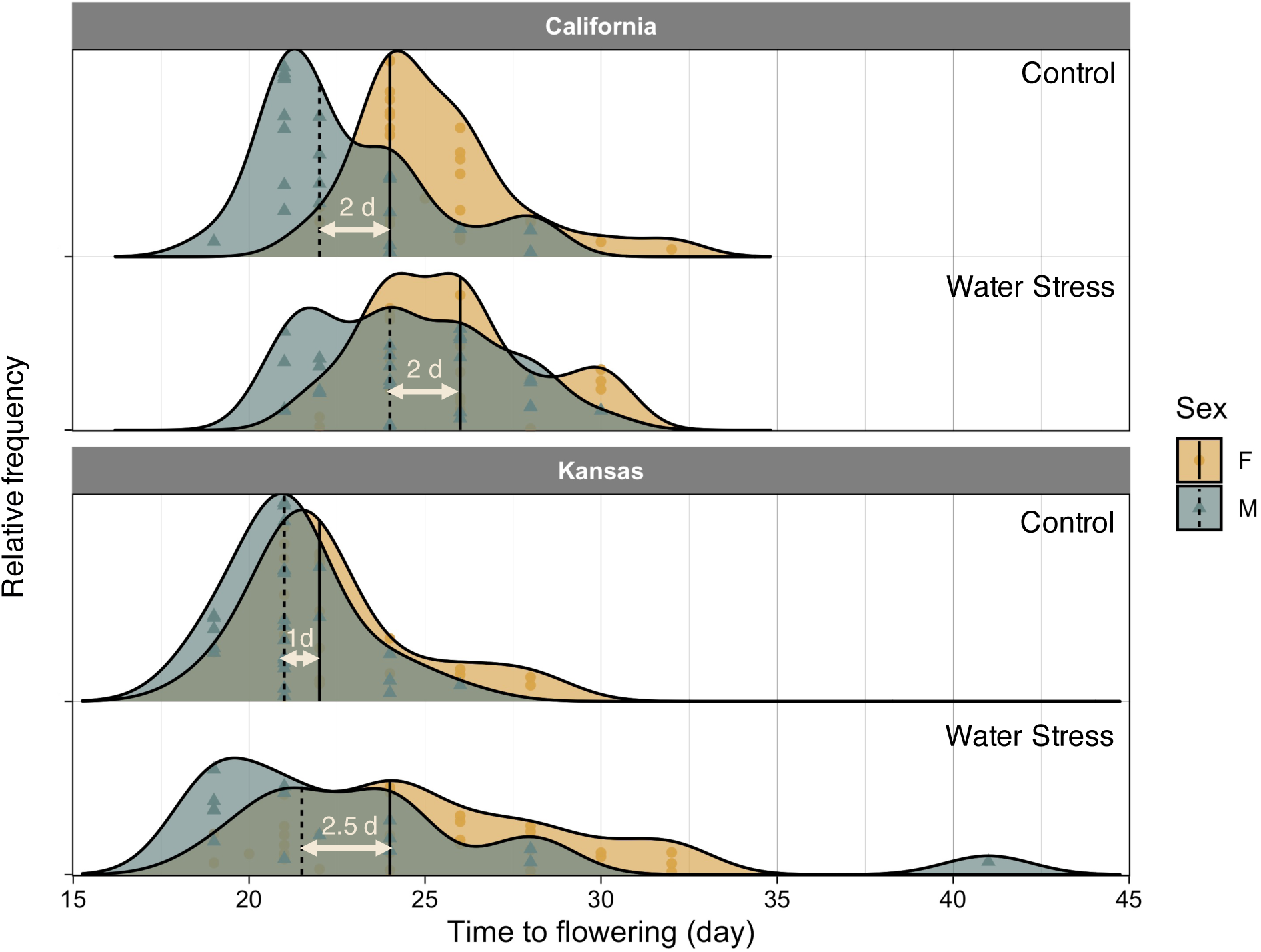
Frequency distribution of time to flowering (emergence of inflorescence) in female (F) and male (M) plants of two *A. palmeri* populations grown under normal watering (control) or water stress conditions. Vertical lines show the location of median time to flowering while symbols indicate observed data. The differences in median flowering time between the two sexes are shown with horizontal arrows.

**Figure 8.**
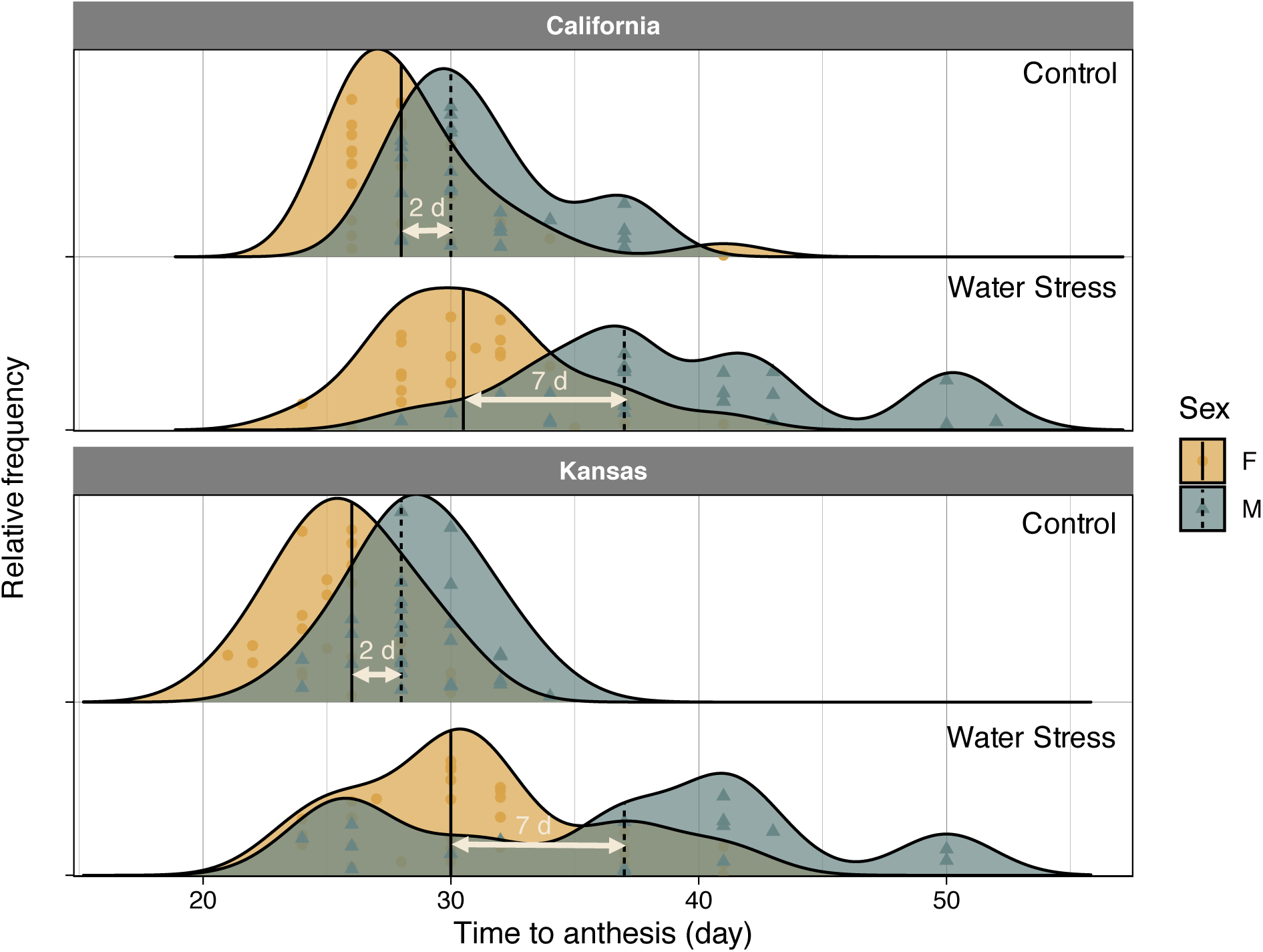
Frequency distribution of time to anthesis in female (F) and male (M) plants of two *A. palmeri* populations grown under normal watering (control) or water stress conditions. Vertical lines show the location of median time to flowering while symbols indicate observed data. The differences in median flowering time between the two sexes are shown with horizontal arrows.

Decreasing the level of irradiance (i.e. greater stress) also postponed *A. palmeri* flowering in the study of Korres et al. (2017). Plants experiencing stressful environments often accelerate their transition to reproductive stage; perhaps as a potential adaptive strategy to produce seed before they die due to stress (Wada and Takeno 2010). However, depending on the intensity and duration of stress and plant development stage, stress may produce an opposite response (i.e. delay the flowering) by slowing the metabolism in plant (Cho et al. 2017). The delay in flowering of *A. palmeri* observed in Korres et al. (2017) and our study might have resulted from a slowed metabolism. However, contrary to our initial hypothesis, the timing of flowering/anthesis in females was less sensitive to stress than males. The prevalence of protandry in dioecious species has been attributed to the greater cost of reproduction incurred by females (Lloyd and Webb 1977; Forest, 2014). As females need to produce seeds, they may undergo an extended period of resources accumulation to ensure sufficient assimilates will be available for allocation to seeds (Lloyd and Webb 1977; Purrington and Schmitt 1998). As a result, stress was proposed to exert a stronger effect on the flowering phenology of females than that of males; a hypothesis that was not supported by our data.

### Implications for weed management

Our study showed that sex can be labile in *A. palmeri* and change in response to external environmental conditions (here water stress). Sex lability can be exploited as new tool for long-term management of *A. palmeri* (and other dioecious weeds): by shifting the sex ratio towards the extremes we might be able to reduce seed production and hence suppress the population growth. A variety of external factors can be used to shift sex towards the desirable direction (e.g. plant hormones, nutrient and water management, light intensity, and herbicides) but currently we have no information about the effects of these factors on the sex expression of *A. palmeri*. Increasing the number of males might offer a more viable and safer management option as males do not contribute to seed production. However, if the population is severely pollen limited, increase in the share of females may also reduce the seed production capacity of population. A quantitative understanding of the relation between thee sex ratio and seed output is needed to determine the proper sex conversion strategy that can potentially disrupt seed production. Our study clearly demonstrated that water stress can result in temporal separation of the two sexes in *A. palmeri*, offering a potential means for ecological management of this weed and perhaps other dioecious weeds. Asynchrony in timing of anthesis can reduce the chance of cross pollination and hence reduce seed production in unisexual weeds such as *A. palmeri* and common waterhemp (*A. tuberculatus* (Moq.) J. D. Sauer). While the direct (negative) effect of water limitation on seed production is well understood, our study suggests a novel mechanism through which water stress (and perhaps other stressors) can reduce seed production indirectly by exacerbating the flowering/anthesis mismatch between the sexes in a dioecious weed. Future work should attempt to disentangle the direct effects of environmental stressors form their indirect effects on seed production, a subject which was out of the scope of the current study.

## Acknowledgments

This research was funded by UC Davis New Research Initiatives and Interdisciplinary Research Grant. The authors would also like to thank Hannah Clifton and William Werner for greenhouse maintenance.

## References

Baker HG (1955) Self-compatibility and establishment after ‘“long-distance” dispersal. Evolution 9:347–349.

Bravo W, Leon RG, Ferrell JA, Mulvaney MJ, Wood CW (2017) Differentiation of life-history traits among Palmer amaranth populations (*Amaranthus palmeri*) and its relation to cropping systems and glyphosate sensitivity. Weed Sci 65: 339–349.

Chailakhyan MK (1979) Genetic and hormonal regulation of growth, flowering, and sex expression in plants. Am J Bot 66: 717–736.

Charlesworth D (1999) Theories of the evolution of dioecy. In Gender and sexual dimorphism in flowering plants (pp. 33–60). Springer, Berlin, Heidelberg.

Charnov EL (1982) The Theory of Sex Allocation. Princeton University Press, Princeton, NJ.

Cho LH, Yoon J, An G (2017) The control of flowering time by environmental factors. Plant J 90:708–719.

Delph LF (1999) Sexual dimorphism in life history. *In* Gender and Sexual Dimorphism in Flowering Plants (Geber, M. A., Dawson, T. E. & Delph, L. F. eds.), pp. 149–173, Springer-Verlag, New York.

Dorken ME, Barrett SC (2004) Sex determination and the evolution of dioecy from monoecy in *Sagittaria latifolia* (Alismataceae). Proc R Soc Lond [Biol] 271: 213–219.

Forero-Montaña J, Zimmerman JK (2010) Sexual dimorphism in the timing of flowering in two dioecious trees in a subtropical wet forest, Puerto Rico. Caribbean J Sci 46: 88–95.

Forrest JR (2014) Plant size, sexual selection, and the evolution of protandry in dioecious plants. Am Nat 184:338–351.

Fox J, Weisberg S, Adler D, Bates D, Baud-Bovy G, Ellison S, Firth D, Friendly M, Gorjanc G, Graves S, Heiberger R (2019) Package ‘car’: Companion to applied regression. https://CRAN.R-project.org/package=car

Freeman DC, Harper KT, Charnov EL (1980) Sex change in plants: old and new observations and new hypotheses. Oecol 47:222–232.

Freeman DC, Vitale JJ (1985) The influence of environment on the sex ratio and fitness of spinach. Botanical Gaz 146:137–142.

Fuller HJ, Ritchie DD (1967) General Botany, Barnes and Noble. Inc., New York.

Geber MA, Dawson TE (1999) Gender and sexual dimorphism in flowering plants. Springer Science & Business Media.

Giacomini D, Westra P, Ward SM (2014). Impact of genetic background in fitness cost studies: an example from glyphosate-resistant Palmer amaranth. Weed Sci 62:29–37.

Heap I (2018) The International Survey of Herbicide Resistant Weeds. Online. Internet Available at: www.weedscience.com (accessed Tuesday, June 17, 2019). https://CRAN.R-project.org/package=car

Juvany M, Munné-Bosch S (2015) Sex-related differences in stress tolerance in dioecious plants: a critical appraisal in a physiological context. J Exp Bot 66:6083–6092.

Keeley PE, Carter CH, Thullen RJ (1987) Influence of planting date on growth of Palmer amaranth (*Amaranthus palmeri*). Weed Sci 35:199–204.

Khryanin VN (2002). Role of phytohormones in sex differentiation in plants. Russian J Plant Physiol 49:545–551.

Korpelainen H (1998) Labile sex expression in plants. Biological Rev 73:157–180.

Korres NE, Norsworthy JK (2017) Palmer amaranth (*Amaranthus palmeri*) demographic and biological characteristics in wide-row soybean Weed Sci 65:491–503.

Korres NE, Norsworthy JK, FitzSimons T, Roberts TL, Oosterhuis DM (2017) Differential response of Palmer amaranth (*Amaranthus palmeri*) gender to abiotic stress. Weed Sci 65:213–227.

Lemen C (1980) Allocation of reproductive effort to the male and female strategies in wind-pollinated plants. Oecol 45:156–159.

Lloyd D G, Webb CJ (1977) Secondary sex characters in plants. Bot Rev 43:177–216.

Lovett Doust J, Cavers PB (1982) Sex and gender dynamics in jack-in-the-pulpit, *Arisaetna triphyllum* (Araceae) Ecology 63:797–808.

Maynard Smith J (1978). The evolution of sex (No. 574.1 S5). Cambridge: Cambridge University Press.

Meagher TR (1988). Sex determination in plants. In Plant reproductive ecology: patterns and strategies. (Lovett Doust, J. & Lovett Doust, L. eds), pp. 125–138. Oxford University Press. New York.

Morgan PW, Drew MC (1997) Ethylene and plant responses to stress. Physiol Plantarum, 100:620–630.

Mulcahy DL (1967) Optimal sex ratio in *Silene alba*. Heredity 22:411–423.

Onyekwelu SS, Harper JL (1979) Sex ratio and niche differentiation in spinach (*Spinacia oleracea* L.). Nature 282:609–611.

Policansky D (1981) Sex choice and the size advantage model in jack-in-the-pulpit (*Arisaema triphyllum*). PNAS 78:1306–1308.

Pospíšilová J, Synková H, Rulcová J (2000) Cytokinins and water stress. Biologia Plantarum 43:321–328.

Purrington CB, Schmitt J (1998) Consequences of sexually dimorphic timing of emergence and flowering in *Silene latifolia*. J Ecol 86:397–404.

R Core Team (2018) R: A language and environment for statistical computing. R Foundation for Statistical Computing, Vienna, Austria. http://www.R-project.org/.

Radosevich SR, Holt JS, Ghersa CM (2007) Ecology of weeds and invasive plants: relationship to agriculture and natural resource management. John Wiley & Sons.

Razanajatovo M, Maurel N, Dawson W, Essl F, Kreft H, Pergl J, Pyšek P, Weigelt P, Winter M, Van Kleunen M (2016). Plants capable of selfing are more likely to become naturalized. Nat Comm 7:13313

Ritz C, Strebig JC, Ritz MC (2016) Package ‘drc’. Analysis of dose-response curves. http://www.bioassay.dk

Sosnoskie LM, Webster TM, Kichler JM, MacRae, AW, Grey TL, Culpepper AS (2012) Pollen-mediated dispersal of glyphosate-resistance in Palmer amaranth under field conditions Weed Sci 60:366–373.

Sutherland S (2004) What makes a weed a weed: life history traits of native and exotic plants in the USA. Oecol 141:24–39.

Thompson AE (1956) Methods of producing first-generation hybrid seed in spinach. Cornell Univ Agr Exp Sta Mem 336:3–48.

Thompson FL, Eckert CG (2004) Trade-offs between sexual and clonal reproduction in an aquatic plant: experimental manipulations vs. phenotypic correlations. J Evol Biol 17:581–592.

Thomson J D, Barrett SC (1981) Selection for outcrossing, sexual selection, and the evolution of dioecy in plants. Am Nat 118:443–449.

Thomson J D, Brunet J (1990) Hypotheses for the evolution of dioecy in seed plants. Trends Ecol Evol 5:11–16.

Van Wychen L (2017) Survey of the most common and troublesome weeds in grass crops, pasture and turf in the United States and Canada. Weed Science Society of America National Weed Survey Dataset. Available: http://wssa.net/wp-content/uploads/2017-Weed-Survey_Grass-crops.xlsx.

Vega-Frutis R, Macías-Ordóñez R, Guevara R, Fromhage L (2014) Sex change in plants and animals: a unified perspective. J Evol Biol 27:667–675.

Wada KC, Takeno K (2010) Stress-induced flowering. Plant Signal Behav 5:944–947.

Webster TM, Grey TL (2015) Glyphosate-resistant Palmer amaranth (*Amaranthus palmeri*) morphology, growth, and seed production in Georgia. Weed Sci 63:264–272.

Wickham H, Chang W, Henry L, Pedersen TL, Takahashi K, Wilke C, Woo K, Yutani H (2019) Package ‘ggplot2’: Create elegant data visualisations using the grammar of graphics, http://ggplot2.tidyverse.org

Zimmerman JK (1991) Ecological correlates of labile sex expression in the orchid *Catasetum viridiflavum*. Ecology 72:597–608.

